# Genetic labeling of embryonically-born dentate granule neurons in young mice using the *Penk^Cre^* mouse line

**DOI:** 10.1101/2023.11.20.567945

**Authors:** Pierre Mortessagne, Estelle Cartier, Maddalena Balia, Murielle Fèvre, Fiona Corailler, Cyril Herry, Djoher Nora Abrous, Arne Battefeld, Emilie Pacary

## Abstract

The dentate gyrus (DG) of the hippocampus is a mosaic of dentate granule neurons (DGNs) accumulated throughout life. While many studies focused on the morpho-functional properties of adult-born DGNs, much less is known about DGNs generated during development, and in particular those born during embryogenesis. One of the main reasons for this gap is the lack of methods available to specifically label and manipulate embryonically-born DGNs. Here, we have assessed the relevance of the *Penk^Cre^* mouse line as a genetic model to target this embryonically-born population. In young animals, *Penk^Cre^* expression allows to tag neurons in the DG with positional, morphological and electrophysiological properties characteristic of DGNs born during the embryonic period. In addition, Penk^Cre^+ cells in the DG are distributed in both blades along the entire septo-temporal axis. This model thus offers new possibilities to explore the functions of this underexplored population of embryonically-born DGNs.

## INTRODUCTION

In most brain regions, neurons are born during embryogenesis. In contrast, in the dentate gyrus (DG) of the rodent hippocampus, genesis of dentate granule neurons (DGNs) starts during the late embryonic period, peaks around the time of birth, and continues at high levels during the adolescent period to reach moderate and then low levels during adulthood (Altman and Bayer, 1990; Schlessinger et al., 1975). The DG is therefore a heterogeneous brain structure composed of sub-populations of DGNs with different temporal origins, i.e. embryonic, early postnatal, adolescent or adult. In the last two decades, the cellular and functional properties of adult-born DGNs have been extensively studied, mainly because the discovery of adult neurogenesis has opened novel avenues for brain repair (Denoth-Lippuner and Jessberger, 2021). Contrastingly, developmentally-born DGNs and in particular embryonically-born DGNs have not been widely characterized. This early-born population is found in the outer part of the granule cell layer (GCL) while later born DGNs subsequently fill the inner part of the GCL in an outside-in fashion, according to their birthdate (Angevine, 1965; Mathews et al., 2010; Schlessinger et al., 1975). In addition, embryonically-born DGNs exhibit distinctive dendritic morphology (Kerloch et al., 2019), display lower excitability than later born DGNs (Laplagne et al., 2007; Save et al., 2018) and do not undergo cell death (Ciric et al., 2019). Although these characteristics suggest that embryonically-born DGNs might have different functional properties, the specific functions and contribution of these neurons to DG functions remain underexplored. A major roadblock to address this question has been the limitation of available methods for specific lineage tracing and genetic manipulation of embryonically-born DGNs.

So far, embryonically-born DGNs have been tagged using tools similar to those described for studying adult-born DGNs such as the nucleotide analogs BrdU or EdU (Mathews et al., 2010; Muramatsu et al., 2007) or retroviruses (Laplagne et al., 2007; Mathews et al., 2010). However, the former approach does not allow for genetic manipulation or analysis of morphology and physiological properties of labeled cells while the latter unspecifically tags all the neurons born during embryogenesis. Other strategies to target embryonically-born DGNs such as *in utero* electroporation of the dentate neuroepithelium (Kerloch et al., 2019) or the transgenic *Ngn2Cre^ERTM^* line (Save et al., 2018) label few embryonic DGNs and/or a large proportion of other types of hippocampal excitatory neurons. Recent work identified an *Osteocalcin-Cre* mouse line (*Ocn-Cre*) as a good tool for specific labeling and manipulating embryonically-born DGNs (Sun et al., 2021). However, with this transgenic mouse, only embryonically-born DGNs in the septal DG are tagged (Sun et al., 2021) excluding the study of embryonically-born DGNs in the temporal DG. This limitation thus further emphasizes the need to develop new tools for the genetic manipulation of the earliest-born DGNs. Two recent studies identified the *Penk* gene, encoding the endogenous opioid polypeptide hormone proenkephalin, in a subpopulation of DGNs with a preferential localization in the outer GCL (Erwin et al., 2020; Habib et al., 2016) characteristic for embryonically-born DGNs. Based on these studies, we hypothesized that *Penk* is a potential candidate gene to identify embryonically-born DGNs.

Here, we aimed to determine whether the *Penk^Cre^* mouse line is a good model to label genetically embryonically-born DGNs. We found that this approach can be relevant in young mice, up to 3 months, but not in older animals.

## MATERIALS AND METHODS

### Animals

*Penk^Cre^ (B6;129S-Penk^tm2(cre)Hze^/J;* RRID:IMSR_JAX:025112) mice and the conditional reporter lines Ai6 (B6.Cg-*Gt(ROSA)26Sor^tm6(CAG-ZsGreen1)Hze^*/J; RRID:IMSR_JAX:007906) (Madisen et al., 2010) and Ai14 (B6.Cg-*Gt(ROSA)26Sor^tm14(CAG-tdTomato)Hze^*/J; RRID:IMSR_JAX:007914) (Madisen et al., 2010) were obtained from The Jackson Laboratory. *Penk^Cre^* heterozygous males were crossed with homozygous Ai6 or Ai14 females to generate *Penk^Cre^;Ai6* or *Penk^Cre^;Ai14* mice. Both males and females were used for the analyses. Mice were maintained under standard conditions (food and water ad libitum; 23+/- 1°C, 12/12 h light-dark cycle; light from 7:00 am to 7:00 pm). These mice were housed, bred, and treated according to the European directive 2010/63/EU and French laws on animal experimentation. All procedures involving animal experimentation and experimental protocols were approved by the Animal Care Committee of Bordeaux (CEEA50) and the French Ministry of Higher Education, Research and Innovation (APAFIS authorization *#*31004).

### Tissue processing

Mice were deeply anesthetized with an intraperitoneal injection of a mixture of pentobarbital (300 mg/kg; Exagon®, Axience) and lidocaine (30 mg/kg, Lidor®, Axience) before transcardial perfusion with ice-cold phosphate buffer saline (PBS, 0.1M, pH=7.3) and then with ice-cold 4% paraformaldehyde (PFA) in PBS. Brains were post-fixed 24h in 4% PFA before being cut coronally and serially (10 series of 40 µm sections or 3 series of 100 µm sections for morphological studies) with a vibratome (Leica).

### EdU injection, detection and quantification

*Penk^Cre^;Ai6* pregnant mice were injected with EdU (BOC Sciences; 10 mg/ml in PBS, i.p., 50 mg/kg body weight) and sacrificed at P35 as illustrated in Figure 1F. Tissue was processed as described above and EdU detection was done in one series of sections using the Click-It^TM^ EdU Alexa Fluor^TM^ 555 Imaging kit (ThermoFischer Scientific) according to the manufacturer’s protocol and was followed by ZsGreen immunostaining. Images were acquired with a Leica SP5 confocal microscope (20 × 0.7NA objective) and cell counts were manually performed using Fiji software (Schindelin et al., 2012). The proportion of ZsGreen^+^ EdU^+^/ZsGreen^+^ among the ZsGreen+ population was quantified in the inner part of the molecular layer and in the GCL.

**Figure 1:**
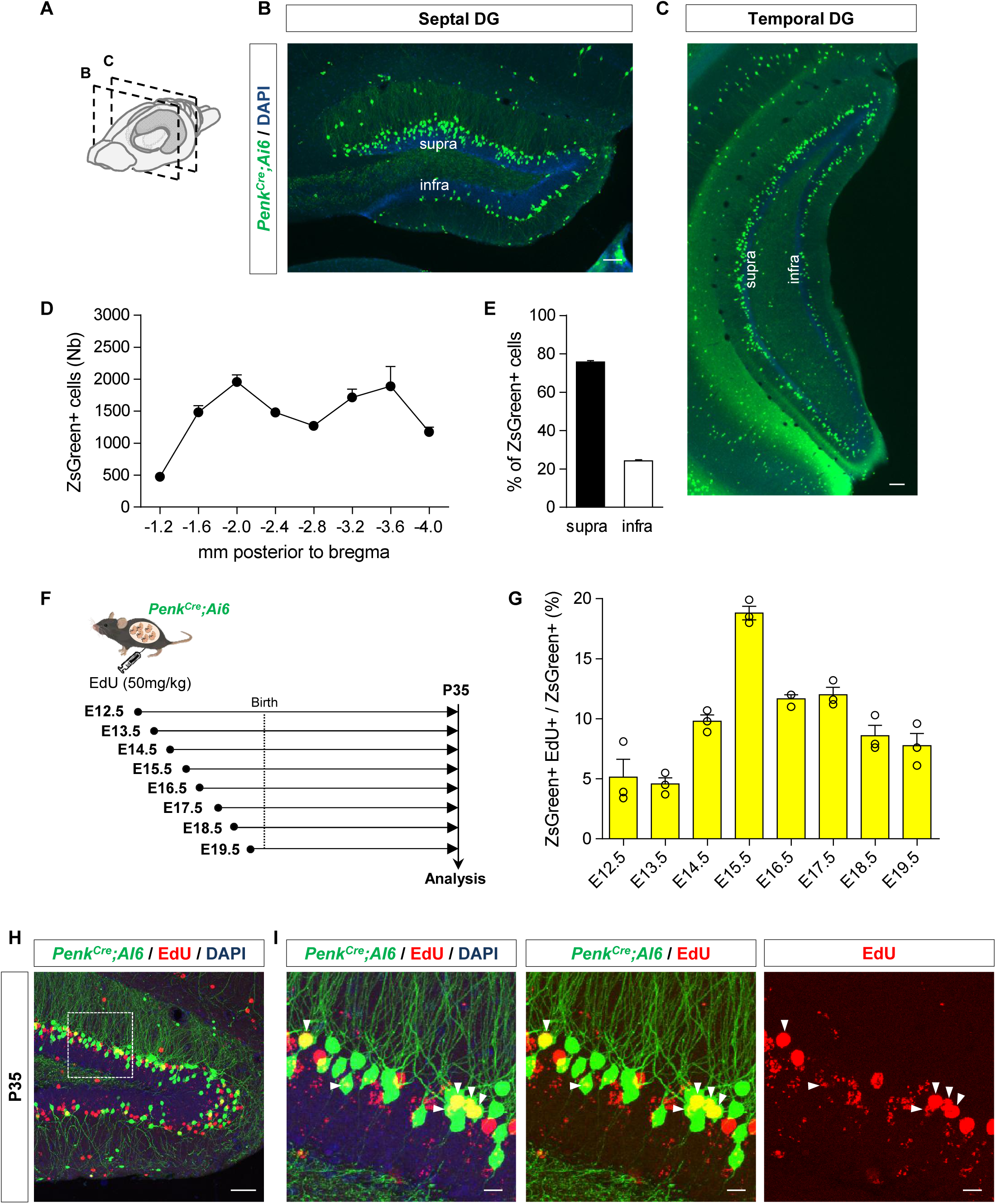
Repartition and birthdating of Penk^Cre^+ cells in the DG. (A) Schematic representation along the mouse brain of the positions of the coronal sections shown in B and C. (B) Septal and (C) temporal DG of *Penk^Cre^; Ai6* mice at P35. (D) Repartition of ZsGreen+ cells along the septo-temporal axis and (E) between the supra- and infra-pyramidal blades of the DG (only cells in the GCL and inner part of the molecular layer were quantified), n = 24 mice. (F) Schematic diagram of the protocol used for birthdating Penk^Cre^+ cells in the DG. (G) Quantification of the percentage of ZsGreen+/EdU cells over total ZsGreen+ cells, n = 3 mice per time point from at least two different litters. (H) Representative image of ZsGreen/EdU/DAPI staining in the DG of *Penk^Cre^;Ai6* mice injected with EdU at E15.5. The white rectangle shows the area enlarged in the insets shown in (I), and white arrowheads indicate double positive cells. Scale bars represent 50 µm (B, C, H) and 10 µm (I).

### Virus injection into the mouse DG

AAV-hSyn-DIO-mCherry (Addgene #50459-AAV8; 2,1.10^13^ vg/ml) was injected bilaterally into the DG of 10-week-old male *Penk^Cre^;Ai6* mice. Thirty minutes before the surgery, mice were injected with Meloxicam (2 mg/kg; Metacam®, Boehringer Ingelheim;). Mice were then anaesthetized with isofluorane in oxygen carrier (TEM SEGA, France) and a cranial subcutaneous injection of 0.1 ml of lidocaine (20 mg/ml; Lidor®, Axience) was performed before settling the mouse into the stereotaxic frame. Betadine was applied, then the skin was cut and a pulled microcapillary glass tube (1-5 μL, Sigma) was placed above bregma. Coordinates of the injection site from bregma were: anteroposterior: −2 mm, mediolateral: −1.8 mm, dorsoventral: −2.1 mm. A small hole was made on the skull using an electric drill, the microcapillary was loaded with the viral solution and introduced within the hole to reach the DG. Then 1 μl of diluted virus (1 in 50) was injected at a rate of 0.5 μl every two minutes and the microcapillary was left in the same position for four minutes after the end of infusion before being removed. Then the skin was stitched with absorbable sutures, betadine and a few drops of lidocaine were applied on the sutures. The mouse was then placed in a recovery chamber (37°C) until it woke up. Mice were sacrificed 2 weeks post-injection and brain tissue processed for morphological analysis (100 µm sections and chromogenic immunohistochemistry).

### Immunohistochemistry

For fluorescent immunohistochemistry, sections were treated with PBS − 0.3% Triton X100 - 3% normal goat serum (NGS) for 45 min after washings in PBS. They were then incubated overnight at 4°C with the following primary antibodies diluted in PBS - 0.3% Triton X100 - 1% NGS: rabbit anti-ZsGreen (1/1000, Clontech, 632474, RRID:AB_2491179) or rabbit anti-DCX (1/2000, Sigma, D9818, RRID:AB_1078699). Sections were then incubated for 2 h at RT with appropriate fluorescent secondary antibodies diluted in PBS – 1% NGS: goat anti-rabbit Alexa Fluor® 488 (1/1000, Invitrogen, A11008, RRID:AB_143165) or goat anti-rabbit Alexa Fluor® 568 (1/1000; Invitrogen, A11036, RRID:AB_10563566). After 2 washes in PBS, cell nuclei were stained by 10 min incubation with DAPI (1/10000, Invitrogen). Sections were mounted using Aqua Polymount (Polyscience). Images were acquired using a Leica SP5 confocal microscope (10× 0.4NA or 40× 1.25NA objective) or using a slide scanner Nanozoomer 2.0HT (Hamamatsu Photonics, France) and a TDI-3CCD camera.

For chromogenic immunohistochemistry, sections were treated with methanol and 0.5% H_2_O_2_ for 30 min after washing in PBS. Sections were washed again in PBS before incubation with a blocking solution containing 3% NGS and 0.3% Triton X100 in PBS for 45 min at room temperature. They were then incubated overnight at 4°C with a rabbit anti-mCherry antibody (1/500, Ozyme, 632496, RRID:AB_10013483) diluted in the blocking buffer. The following day, sections were incubated with a biotin-labeled goat anti-rabbit secondary antibody (1/200, Vector Labs, BA-1000, RRID:AB_2313606) diluted in PBS - 0.3% Triton X100 – 1 % NGS. Immunoreactivities were visualized by the biotin-streptavidin technique (ABC kit; Vector Laboratories; PK-4000) with 3, 3′-diaminobenzidine (DAB) as chromogen.

To perform morphometrical analysis after electrophysiology, biocytin filled single neurons were reacted with streptavidin. After several washes in PBS, sections were blocked with 0.3% Triton X100, 1% bovine serum albumin (BSA) in PBS for 1 h. Thick slices were then incubated overnight at 4°C with streptavidin coupled to Alexa-568 (1/500, Invitrogen, S11226, RRID:AB_2315774) diluted in PBS containing 0.3% Triton X100 and 1% BSA. After 2 washes in PBS, cell nuclei were stained by 10 min incubation with DAPI (1/10000, Invitrogen). Sections were mounted using PVA (Sigma, P8136) - DABCO (Sigma, D2522).

### Neuronal structure tracing and quantification

Dendrites and cell body of mCherry positive DGNs labeled with DAB (Figure 2) or biocytin-filled DGNs (Figure 3) were traced using a 100× 1.4NA objective and a semiautomatic neuron tracing system (Neuron Tracing from Neurolucida® software; MicroBrigthField) as previously described (Kerloch et al., 2019). Only DGNs in the suprapyramidal blade were used as in Kerloch et al. (2019). Cells with primary dendrites cut off in the proximal region (inner third of the molecular layer and below) were excluded as well as cells with more than three dendrites cut off in the molecular layer. Measurements of primary dendrite number, dendritic length, branching angle and relative position in the GCL were performed with Neurolucida software. Primary dendrites are the dendrites emerging from the soma (Figure 2D). The dendritic length is the sum of the length of all dendritic branches (Figure 2E). The angle between the two most lateral branches (branching angle) was measured from the center of the cell body (Figure 2F). The relative position of the cell in the GCL was defined as follows: the inner border of the GCL was defined as the baseline to measure cell position and the perpendicular distance from this baseline to the center of the cell body was measured and normalized by the thickness of the GCL (Figure 2G).

**Figure 2:**
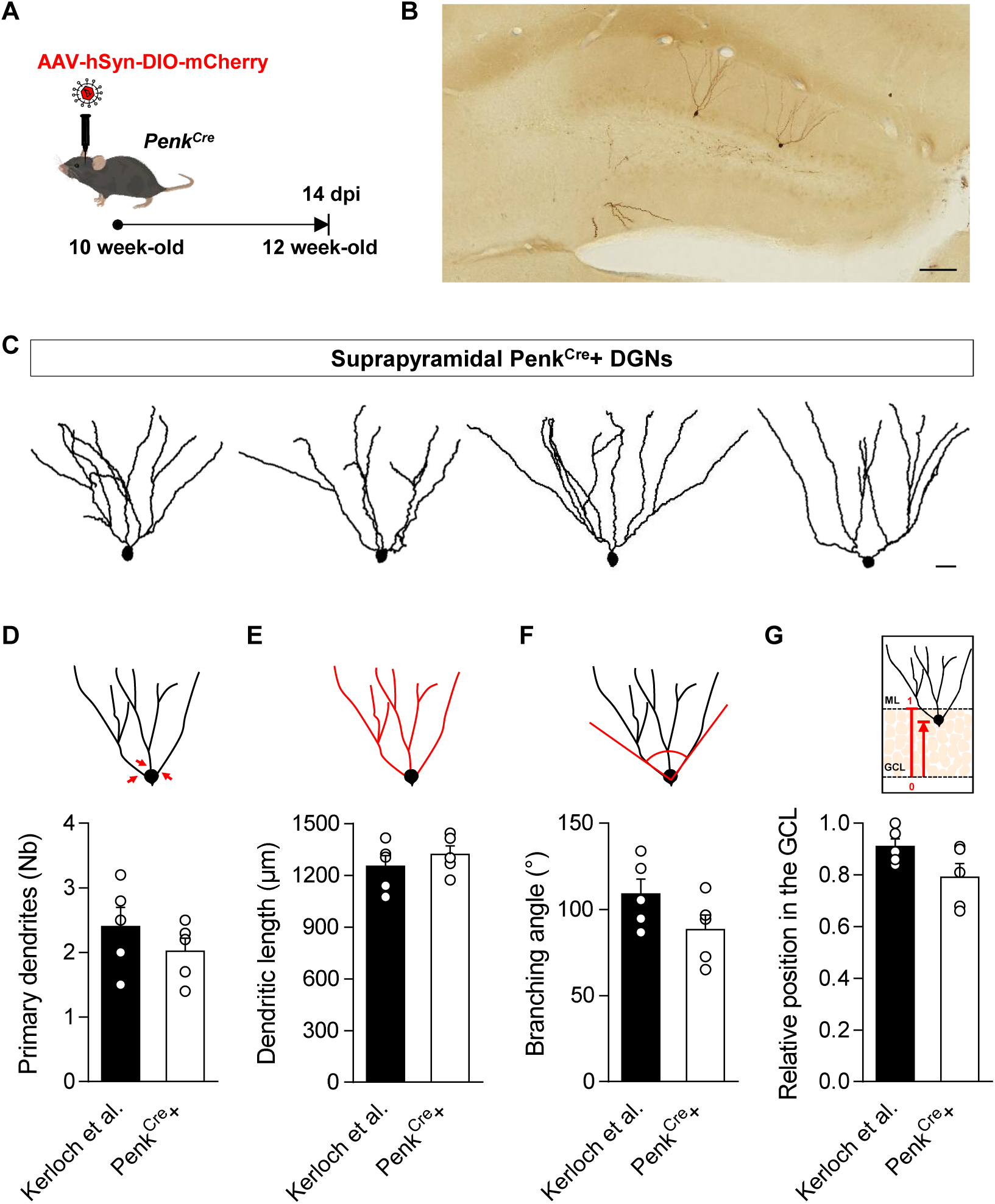
Penk^Cre^+ cells in the DG have a morphology similar to DGNs born at E14.5. (A) Experimental design. (B) Sparse labeled cells after injection of AAV-hSyn-DIO-mCherry virus into the DG of a *Penk^Cr^*^e^ mouse. (C) Representative tracings of Penk^Cre^+ neurons in the DG of 12 week-old mice. Comparison of (D) the number (Nb) of primary apical dendrites, (E) the total dendritic length, (F) the branching angle and (G) the relative position in the GCL of Penk^Cre^+ neurons with DGNs tagged at E14.5 in Kerloch et al. (2019) with in utero electroporation. Penk^Cre^+ neurons were traced from n = 5 mice (30 neurons in total with a minimum of 3 neurons per animal) and were compared to the DGNs from Kerloch et al. (n = 5 mice) using Mann-Whitney test. Scale bars represent 100µm (B) and 20 µm (C).

**Figure 3:**
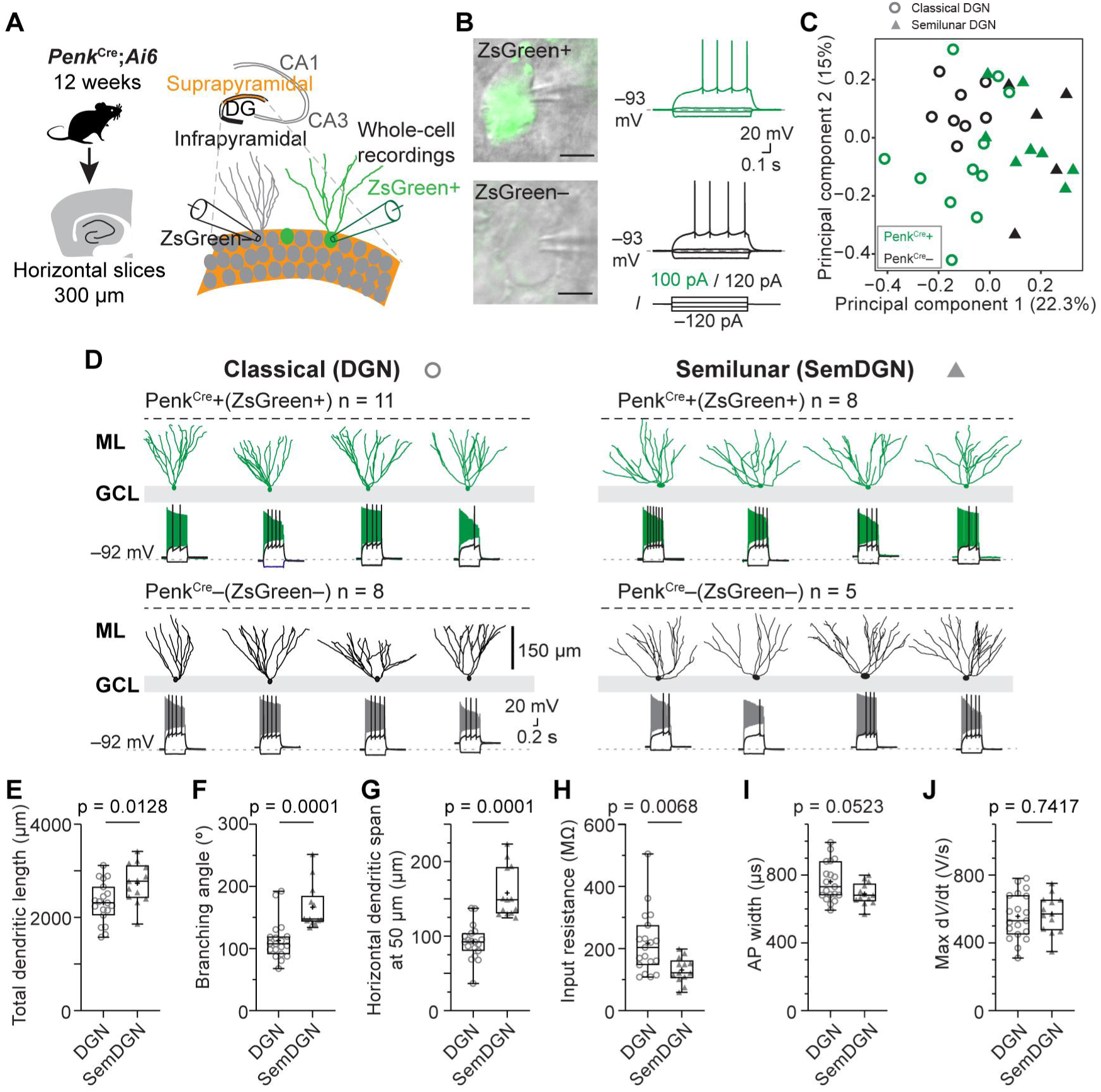
Morpho-electric analysis of Penk^Cre^+ DG neurons reveals a heterogeneous neuronal population similar to outer PenkCre- DGNs. (A) Schematic of the experimental workflow showing the targeting of Penk^Cre^+ and Penk^Cre^- DGNs in the first outer layer of the suprapyramidal blade in horizontal acute slices by whole- cell patch-clamp. (B) Oblique contrast image examples of ZsGreen fluorescent Penk^Cre^+ (top) and non-fluorescent Penk^Cre^- (bottom) DGNs with example voltage responses. (C) Principal component (PC) analysis of morpho-electric properties reveals no clustering of Penk^Cre^+ (green) or Penk^Cre^– (black) DGNs. Circles identify classical DGNs, while closed triangles represent semilunar granule neurons as in D. (D) Morphological reconstructions and corresponding voltage responses of recorded DG neurons. Reconstructions identified two distinct morphological classes: classical DG and semilunar DG neurons. Examples are shown for several *Penk^Cre^*+ and *Penk^Cre^*– neurons. The voltage traces show subthreshold response, the action potential firing at threshold and the maximal action potential frequency response. Scale bars are for all examples shown. (E) (F) (G) (H) (I) (J) Box plots for three morphological parameters and three electrophysiological parameters reveal some differences between classical DGNs (DGN) and semilunar DGNs (semDGN). A Mann-Whitney test was performed for (E), (F) and a t-test was performed for (G), (H), (I), (J) with p-values given on the corresponding graphs. For all box plots, the box represents the 25^th^ and 75^th^ percentile, the line dividing the box the median, the whiskers indicate maximum and minimum and the mean is represented with a cross. All data points are represented. Scale bar represents 5 µm (B).

### Acute brain slice preparation

Acute slices were prepared from male and female *Penk^Cre^;Ai6* mice that were 85 ± 2 (n = 10) days old on the day of the experiment. Mice were deeply anesthetized and subsequently transcardial perfused with ice-cold ACSF (in mM: 100 Sucrose, 60 NaCl, 25 NaHCO_3_, 1.25 NaH_2_PO_4_, 2.5 KCl, 20 Glucose, 1 CaCl_2_, 6 MgCl_2_) saturated with 95% O_2_ and 5% CO_2_. After perfusion, mice were decapitated and the brain was quickly removed and dissected in the same ice-cold cutting solution. 300 µm thick horizontal slices containing the hippocampus were cut with a vibratome (VT1200S, Leica Microsystems, Germany) and collected and incubated in storage ACSF solution (in mM: 125 NaCl, 25 NaHCO_3_, 1.25 NaH_2_PO_4_, 3 KCl, 25 Glucose, 1 CaCl_2_, 6 MgCl_2_) saturated with 95% O_2_ and 5% CO_2_ for 35 min at 35°C. Afterward slices were transferred to room temperature and stored in the same solution for the duration of the experimental day.

### Whole-cell electrophysiology

For electrophysiology experiments, slices were transferred to a submerged chamber under an upright microscope either Olympus BX51WI (Olympus France, Rungis, France) or LN-scope (Luigs-Neumann, Ratingen, Germany) and perfused with heated (33 ± 2°C) ACSF (in mM: 125 NaCl, 25 NaHCO_3_, 1.25 NaH_2_PO_4_, 3 KCl, 25 Glucose, 2 CaCl_2_, 1 MgCl_2_) saturated with 95% O_2_ and 5% CO_2_. DGNs were visualized with oblique contrast using near-infrared (720 nm) LED illumination and a CMOS camera (Blackfly, Teledyne FLIR LLC, Wilsonville, OR, USA). Penk^Cre^ positive DGNs or non-fluorescent control neurons were identified by their expression or lack of ZsGreen using a blue LED (470 nm or 460 nm) and a filter set optimized for GFP that included a long band-pass emission filter (GFP-30LP-B-000, Semrock, Rochester, NY, USA). Fluorescent neurons were targeted in the upper layer of the suprapyramidal blade of the DG and non-fluorescent neurons were targeted in the same layer usually adjacent to fluorescent neurons. Whole-cell patch-clamp recordings of DGNs were made with an intracellular solution containing (in mM) 130 K-Gluconate, 10 KCl, 10 HEPES, 4 Mg-ATP, 0.3 Na_2_-GTP, 10 Na_2_-phosphocreatine and biocytin (5 mg/ml) for post-hoc reconstruction of the morphology. Recording pipettes were pulled on a vertical puller from borosilicate glass capillaries (1.5 mm outer diameter, 0.86 mm inner diameter). When filled with intracellular solution the open tip resistance ranged from 3.6 to 7.7 MΩ. The liquid junction potential (LJP) was estimated to be –12 mV and reported voltage values are corrected for the LJP. All voltage-clamp and current-clamp recordings were made with Sutter IPA2 amplifiers (Sutter Instrument, Novato, CA, USA) controlled by SutterPatch software (Sutter). Recordings were filtered at 10 kHz (Bessel) and sampled with a minimum frequency of 20 kHz. At the end of recording, the slice was removed from the recording chamber and fixed in fresh 4% PFA in PBS during 12 hours and subsequently washed with PBS.

### Analysis of electrophysiology recordings

Analysis of recordings was performed with standardized analysis protocols in SutterPatch or after exporting data to Axograph (Axograph.com). Input resistance was determined from linear fits to voltage responses in current clamp around the resting membrane potential. Resting membrane potential was determined in current-clamp from segments when I = 0 pA. Synaptic events were analyzed with Axograph X using a template matching detection algorithm. For this, recordings were low-pass filtered at 1 kHz and subsequently events were automatically detected. Detection threshold was set to minimally 2.5 times SD and a cut-off was made for all events smaller than –3 pA. Action potentials (APs) were automatically offline analyzed with the AP analysis module in SutterPatch and a threshold for detection of 0 mV. From these automatically detected events we extracted the threshold (when voltage change >10 V/s), the AP amplitude (threshold to peak) the maximum rise (maximum dV/dt), the AP width at 50% of the peak amplitude and the AP frequency. Data was plotted in Prism 9 for visualization. Principal component analysis and display of morpho-electric parameters (5 morphological and 12 electric parameters) was performed in R using the *prcomp* command.

### Statistical analysis

For Figures 1, 2, 4 and S2, data points shown are means from biological replicates that were computed from technical replicates as specified in the legends. Statistical analysis was performed from the biological replicates which are shown in the respective figures. For all data from electrophysiology experiments, we tested for normal distribution using a Shapiro-Wilk test. Normal distributed data was tested with an unpaired t-test, while non-normal distributed data were tested with a non-parametric Mann-Whitney test. All statistical analyses were performed with GraphPad Prism (v9) software. Results are presented as mean ± s.e.m (standard error of the mean) in Figures1, 2, 4 and S2. Statistical tests and sample size (n) are indicated in Figure legends. Statistical values are mentioned in the text, legends, figures or table.

## RESULTS

### Penk^Cre^+ cells in the DG are largely born during embryogenesis

As a first step to characterize the *Penk^Cre^* mouse, we crossed this line with the Ai6 reporter, expressing bright ZsGreen fluorescence following Cre-mediated recombination (Madisen et al., 2010), and sacrificed the animals at postnatal day 35 (P35). This time point was chosen to compare our data with the labeling obtained with the *Ocn-Cre;Ai9* (TdTomato) mice (Sun et al., 2021). As expected, ZsGreen+ cells were observed in the DG (Figures 1 A-C) but also in other parts of the brain such as in layers 2 and 6 of the motor and somatosensory neocortex and the striatum as already reported (Daigle et al., 2018) as well as in piriform cortex, the lateral entorhinal cortex, suprachiasmatic nucleus and interpeduncular nucleus (Figure S1). Interestingly, in the DG, labeled cells were distributed along the septo-temporal axis (Figures 1 A-D) and most of them (75.8 ± 0.6%) were detected in the suprapyramidal blade (Figure 1E). More importantly, these cells were mainly located at the border between the GCL and inner molecular layer (Figures 1B-C), a position reminiscent of embryonically-born DGNs. These results thus reinforced our hypothesis that Penk^Cre^+ cells in the DG could be embryonically-born DGNs.

Next, to determine the exact time-point of DGN birth during embryogenesis, we used EdU for birth dating. As shown in Figure 1F, EdU was i.p. injected into pregnant *Penk^Cre^;Ai6* mice at different time points. Mice were then analysed at P35 and the proportion of ZsGreen+ cells expressing EdU was quantified (Figures 1G-I). Using this strategy, we found that Penk^Cre^+ cells in the DG are mainly generated during late embryogenesis with a peak of birth at E15.5. Interestingly, no major differences were observed between the suprapyramidal and infrapyramidal blades (Figure S2). Thus, the majority of fluorescent Penk^Cre^+ cells in both blades of the DG are in majority born during embryogenesis.

### Penk^Cre^+ cells have position and morphological characteristics of embryonically-born DGNs

To further characterize Penk^Cre^+ cells in the DG, we studied their morphology. For morphological reconstructions, we applied a sparse labeling strategy. We injected AAV-hSyn-DIO-mCherry into the DG of 10-week old *Penk^Cre^* mice and analyzed the morphology of mCherry+ cells (= Penk^Cre^+) 14 days later, i.e. when the animals were 12 week-old (Figure 2A, B). We compared the morphology of these cells with that of 12 week-old embryonically-born DGNs that were labeled by in utero electroporation at E14.5 from our previous study (Kerloch et al., 2019). For proper comparison, we studied only DGNs in the suprapyramidal blade and included both classical DGNs and semilunar granule neurons as done in Kerloch et al. (2019) (Figure 2C). We found that Penk^Cre^+ cells have a similar number of primary dendrites (Figures 2 C, D; t_8_ = 1.05, p = 0.32), dendritic length (Figures 2 C, E; t_8_ = 0.86, p = 0.42) and branching angle (Figures 2 C, F; t_8_ = 1.68, p = 0.13) as DGNs born at E14.5. In addition, we confirmed that Penk^Cre^+ cells are mainly located in the outer GCL, as shown by their relative position in the GCL (0.79±0.05 for Penk^Cre^+ cells, t_8_ = 1.94, p = 0.09) (Figure 2G). Based on their location in the GCL and their morphology, Penk^Cre^+ DGNs correspond to embryonically-born DGNs.

### Penk^Cre^+ DGNs have morpho-electrical properties similar to neighboring DGNs

We next examined the electrophysiological profile of Penk^Cre^+ DGNs to determine whether these cells exhibit distinct electrical characteristics and thus represent a discrete neuronal population. We whole-cell recorded from Penk^Cre^+ DGNs in acute *ex vivo* hippocampal brain slices in the outer part of the suprapyramidal blade of 12 week-old mice (*Penk^Cre^;Ai6,* Figure 3A and S3A) and compared them to non-fluorescent neighboring DGNs (Figure 3B). We combined these recordings with biocytin-based post-hoc morphological reconstructions. Physiological properties such as resting membrane potential, input resistance and sag-ratio, as well as post-synaptic currents were similar when comparing Penk^Cre^+ and Penk^Cre^– neurons (Table 1, Figure 3B and S3B-D). Similarly, action potential (AP) properties and single-neuron morphological parameters were comparable between Penk^Cre^+ and Penk^Cre^– DGNs (Table 1). We additionally performed a principal component analysis of electro-morphological properties for unbiased analysis that also showed no separation between these two populations (Figure 3C). These data suggest that Penk^Cre^+ DGNs are indistinguishable from potential embryonically-born DGNs that are Penk^Cre^–.

**Table 1.** – Electrophysiological and morphological properties of Penk^Cre^+ and Penk^Cre^- neurons located in the suprapyramidal blade of the DG. All measurements were obtained at an age of 12 ± 0.2 weeks from whole-cell patch-clamp recordings and subsequent single neuron reconstructions.

| Parameter | Penk<sup>Cre+</sup> (Zs-Green<sup>+</sup>) | Penk<sup>Cre-</sup> (Zs-Green<sup>-</sup>) | Statistical test | P value |
| --- | --- | --- | --- | --- |
| Resting membrane potential (mV) | $-89.8 \pm 0.8$ (n = 19) | $-91.0 \pm 1.6$ (n = 13) | Mann-Whitney | 0.25 |
| Input resistance (MΩ) | $191 \pm 24$ (n = 19) | $169 \pm 21$ (n = 13) | Mann-Whitney | 0.70 |
| Sag ratio (%) | $0.83 \pm 0.1$ (n = 19) | $0.68 \pm 0.1$ (n = 13) | Unpaired t-test | 0.38 |
| PSC frequency (Hz) | $0.68 \pm 0.22$ (n = 19) | $0.66 \pm 0.08$ (n = 12) | Mann-Whitney | 0.22 |
| PSC amplitude (pA) | $-8.1 \pm 0.6$ (n = 19) | $-6.7 \pm 0.4$ (n = 12) | Unpaired t-test | 0.10 |
| **Long current step (800 ms) single action potential (AP) properties** |  |  |  |  |
| AP threshold (mV) | $-48.2 \pm 0.7$ (n = 19) | $-48.2 \pm 1.5$ (n = 13) | Unpaired t-test | 0.97 |
| AP current threshold (pA) | $113 \pm 10$ (n = 19) | $123 \pm 14$ (n = 13) | Mann-Whitney | 0.46 |
| AP amplitude (mV) | $90.4 \pm 1.3$ (n = 19) | $90.8 \pm 1.7$ (n = 13) | Unpaired t-test | 0.84 |
| AP rate of rise (max dV/dt) | $533 \pm 29$ (n = 19) | $606 \pm 31$ (n = 13) | Unpaired t-test | 0.11 |
| AP width (μs) | $740 \pm 26$ (n = 19) | $722 \pm 24$ (n = 13) | Mann-Whitney | 0.97 |
| AP frequency at threshold potential (Hz) | $3.9 \pm 0.4$ (n = 19) | $4.2 \pm 0.4$ (n = 13) | Mann-Whitney | 0.59 |
| AP 1<sup>st</sup> spike latency at threshold potential (s) | $0.32 \pm 0.05$ (n = 19) | $0.30 \pm 0.05$ (n = 13) | Mann-Whitney | 0.54 |
| First Inter-spike-interval (ISI) at threshold (Hz) | $5.5 \pm 0.5$ (n = 19) | $14.6 \pm 8.1$ (n = 13) | Mann-Whitney | 0.10 |
| Average Maximum firing rate (Hz) | $38.7 \pm 2.6$ (n = 19) | $40.1 \pm 1.7$ (n = 13) | Unpaired t-test | 0.69 |
| **Morphological parameters** |  |  |  |  |
| Total dendritic length (μm) | $2410 \pm 102$ (n = 19) | $2597 \pm 141$ (n = 13) | Unpaired t-test | 0.28 |
| Branching angle ($^{\circ}$) | $140 \pm 10$ (n = 19) | $127 \pm 11$ (n = 13) | Unpaired t-test | 0.39 |
| Number primary dendrites | $3.4 \pm 0.3$ (n = 19) | $3.2 \pm 0.3$ (n = 13) | Mann-Whitney | 0.84 |
| Relative position GCL | $0.95 \pm 0.02$ (n = 19) | $0.93 \pm 0.02$ (n = 13) | Mann-Whitney | 0.47 |
| Dendritic span at 50 μm (μm) | $123 \pm 11$ (n = 19) | $113 \pm 10$ (n = 13) | Unpaired t-test | 0.54 |

In addition, the morphological reconstructions unmasked the two morphologically distinct embryonically-born DGN populations, i.e. classical DGNs and semilunar granule neurons (Figure 3D). These two neuron types were previously found to differ in their intrinsic properties (Williams et al., 2007) and semilunar granule neurons have been shown to express *Penk* (Erwin et al., 2020). Interestingly, the principal component analysis almost completely separated classical DGNs (circles) and semilunar granule neurons (closed triangles) with only few overlapping cells (Figure 3C and S3E). We regrouped and pooled neurons by morphological category and compared morpho-electric parameters that were identified to contribute most to the first two principal components (Figure S3E). In agreement with a larger dendritic tree and wider dendritic spread of semilunar granule cells (Figure 3E-G) their input resistance was also lower (Figure 3H), but AP properties were similar between semilunar granule cells and DGNs (Figure 3I-J).

In summary, our data suggests that Penk^Cre^+ DGNs cannot be distinguished in the outer GCL by their electrical properties but encompasse both classical DGNs and semilunar granule neurons, each characterized by distinct morpho-electric properties.

### The number of Penk^Cre^+ cells in the DG increases during young adulthood

Since late embryonically-born DGNs have been reported to persist in rats at least up to 6 months (Ciric et al., 2019), we then investigated whether the Penk-Cre+ population was also stable throughout young adulthood. We analysed brains of *Penk^Cre^;Ai6* mice at 1, 3 and 6 months and determined the prevalence of Penk-Cre+ neurons and observed an increased number of ZsGreen+ cells in the DG with age (Figure 4A and S4). ZsGreen+ cells progressively filled the inner part of the GCL in an outside-in fashion (Figures 4B-C). Nonetheless, it should be noted that the majority of labeled cells were found in the outer GCL regardless of the age of the animals but this proportion decreases with age (Figure 4C). We confirmed this observation by crossing *Penk^Cre^* mice with the Ai14 reporter mouse line. Tdtomato expression was observed in a similar pattern, thus excluding the possibility that the observed effect was due to the fluorescent reporter (Figure S5A). These data indicate that all populations of DGNs (born from embryogenesis to adulthood) express *Penk* at least at one time point during their life. Moreover, since the increase became obvious between 3 and 6 months, this suggests that *Penk* is expressed at a late mature stage during DGN development. This was further supported by RNA in situ hybridization for *Penk* from the Allen Brain Atlas showing almost no expression at P14 and then a progressive expression from the outer GCL to the inner GCL with age (Figure S5B). In addition, no ZsGreen+ cells were found positive for doublecortin (DCX), a marker of immature neurons, in the DG of 1, 3 or 6 month-old mice (Figure 4D) (co-localization was assessed from 3 animals at each time point). Altogether, these results demonstrate that the *Penk^Cre^* mice allow to label embryonically-born DGNs when limiting analysis to young stages, up to 3 months old.

**Figure 4:**
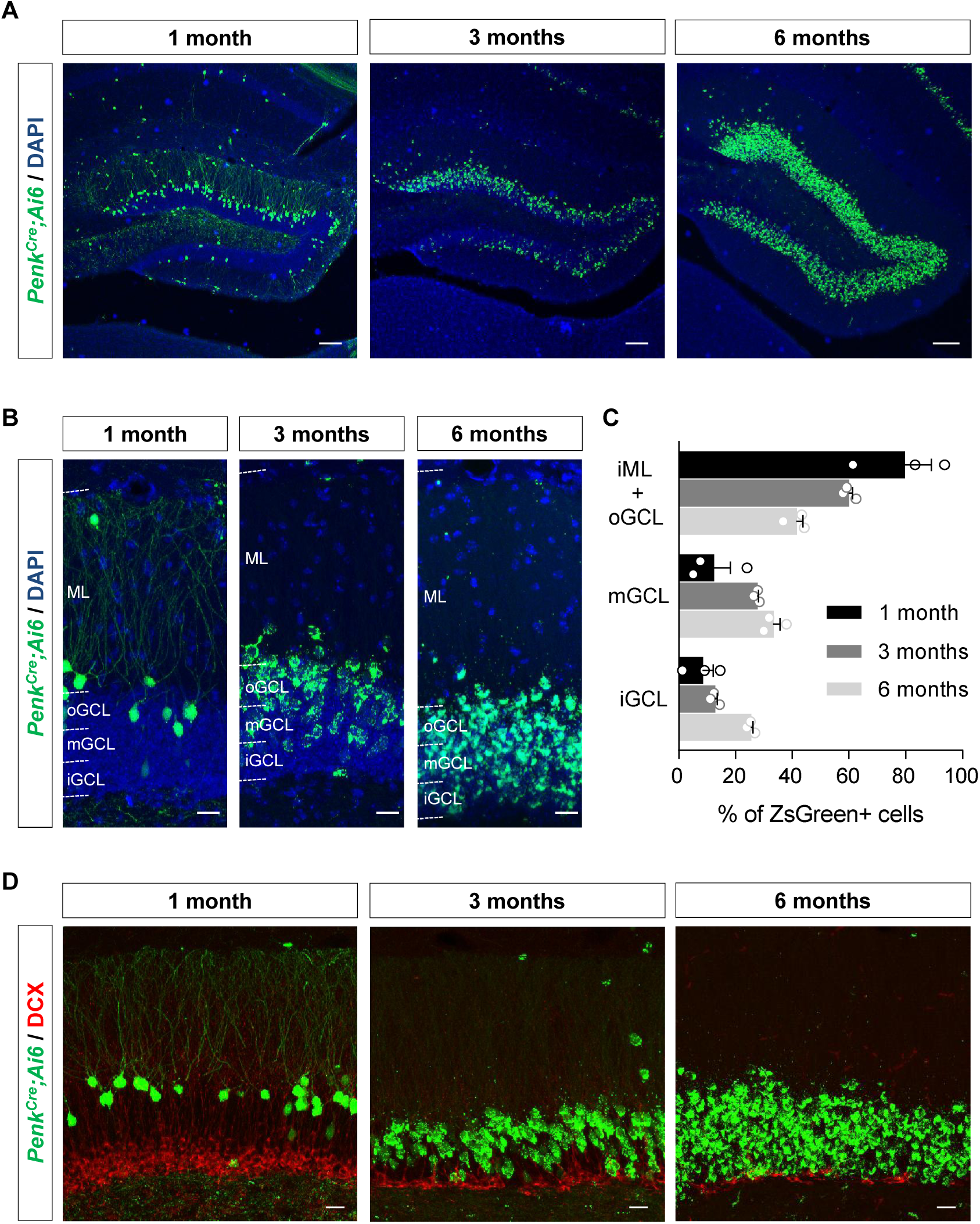
The number of Penk^Cre^+ cells in the DG increase during young adulthood. (A) Septal DG of *Penk^Cre^; Ai6* mice at 1, 3 and 6 months. (B) Images illustrating the repartition of ZsGreen+ cells in the inner GCL (iGCL), middle GCL (mGCL), outer GCL (oGCL) and molecular layer (ML) at the different time points. (C) The quantification graph shows the distribution of ZsGreen+ cells in the different domains at the three ages. iML: inner ML. n = 3 animals per time point. (D) DCX staining in the DG of *Penk^Cre^; Ai6* mice at 1, 3 and 6 months. Scale bars represent 100 µm (A) and 20 µm (B, D).

## DISCUSSION

To date, only a few tools exist to genetically label and manipulate embryonically-born DGNs, thus limiting our understanding of their functions in the brain. In this study, we have assessed the relevance of the *Penk^Cre^*knock-in mouse line as a genetic model to specifically target this population of DGNs. Our data indicate that this model is relevant in young animals, up to 3 months, when studying embryonic born neurons in the DG. Indeed, until 3 months, tagged cells in the outer GCL of the DG have positional, morphological and electrophysiological properties typical of embryonically-born DGNs. At time points beyond 3 months, all DGNs express the fluorescent reporter, thus restricting the use of *Penk^Cre^* mice to young stages when addressing questions that consider the birthdate of the DGNs. For these questions, the used *Penk^Cre^* line has limitations that need to be taken into account, and future advancements should aim to develop complementary tools to overcome this drawback.

### Repartition and proportion of Penk^Cre^+ DGNs in the DG

We found that Penk^Cre^+ cells are distributed along the entire septo-temporal axis allowing to explore the functional distinctions between septal and temporal embryonically-born DGNs. Being able to target neurons on the entire septo-temporal axis of the DG is particularly significant in light of the prevailing view that associates the septal hippocampus with memory and spatial navigation, while attributing anxiety-related behaviors to the temporal hippocampus (Bannerman et al., 2004). Importantly, this provides an advantage to the *Penk^Cre^* model over the previously published *Ocn-Cre* mouse line, which restricts studies exclusively to septal embryonically-born DGNs (Sun et al., 2021).

While our data indicate a preferential localization of Penk^Cre^+ DGNs in the suprapyramidal blade of the DG in young animals, analysis of 6-month-old mice showed a less restrictive expression of the reporter. This contrast might stem from differential development of the two blades. The suprapyramidal blade is formed before the infrapyramidal one (Frotscher et al., 2007) and *Penk* driven Cre expression could thus indicate distinct maturation stage of supra and infra DGNs (Pompeiano and Colonnese, 2023). Moreover, the subsequent expression of *Penk* could also reflect divergent functions between the DGNs of the two blades (Berdugo-Vega et al., 2023). For example, neurons of the suprapyramidal blade are selectively recruited during exposure to a novel environment (Chawla et al., 2005) and these activated DGNs express *Penk* (Erwin et al., 2020).

In the adult mouse DG, *Penk*-expressing DGNs were estimated to account for 5% of the total DGN population, representing a rare subtype of DGNs in mice (Erwin et al., 2020). However, our data suggest that this percentage might depend on the age of the animals. Hence, in young animals, only DGNs generated during embryogenesis might be sufficiently mature to express *Penk* while in 6 month-old animals most DGNs have reached maturity and express *Penk*. Nonetheless, our data do not rule out the possibility that the small proportion observed by Erwin et al. is linked to a transient expression of *Penk* following neuronal activation and that the reporter mice used show the temporal summation of *Penk* expression over their lifespan, which is well visible in 6 months old mice.

### Open questions regarding proenkephalin function in DGNs

While we used *Penk^Cre^* driven reporter expression to identify a subpopulation of DGNs, we cannot ignore the expression of proenkephalin and the subsequent release of the signaling enkephalin peptides from these DGNs. These different cleaved neuropeptides activate opioid receptors (mu-, delta-, or kappa-) and regulate several physiological functions, including pain perception, aggressive-behavior, but also responses to stress (Fricker et al., 2020; Kieffer and Gaveriaux-Ruff, 2002; Konig et al., 1996). Notably, *Penk* is specifically upregulated in DG engram neurons after fear conditioning suggesting an important role in memory consolidation (Rao-Ruiz et al., 2019). However, the specific contribution of DGN-derived proenkephalin to these various functions remains unknown.

Additionally, the identity of the downstream targets that are modulated by the enkephalins derived from *Penk*-expressing DGNs still have to be identified to clarify the function of this neuropeptide precursor. Interestingly, in a scRNA-seq dataset obtained from the mouse DG (Habib et al., 2016), one of the enkephalin receptors *opioid receptor delta 1* (*Oprd1*) was found to be exclusively expressed in GABAergic subclusters and to overlap largely with *parvalbumin* (*Pvalb*) suggesting that Penk release from DGNs could modulate interneuron activity within the DG. Other possible targets are neurons in the hippocampal CA2/CA3 region where DGNs project their axons. Overall, further functional *in vivo* experiments are needed to decipher the contribution of enkephalin co-release from DGNs, which might help to understand the role of embryonically-born DGNs in learning paradigms especially in young animals.

In conclusion, the *Penk^Cre^*mouse line represents a new and complementary tool to study embryonically-born DGNs in young mice and our characterization will help to design reliable experiments and accurate interpretation of future experiments.

## ACKNOWLEDGMENTS

This work was supported by INSERM, ANR (ANR-19-CE16-0014-01 and ANR-22-CE16- 0026-02 to E.P.), FRM (Fondation pour la Recherche Medicale, EQU202203014657 to D.N.A.), a Neurocampus Startup grant provided by the Region Aquitaine (2018.599 to A.B.), GPR Brain_2030 (PhD extension grant to P.M.), IDEX ATTRACTIVITE Chaires Neurocampus of the University of Bordeaux (AB) and financial support from the French government in the framework of the University of Bordeaux’s IdEx “Investments for the Future” program / GPR BRAIN_2030. We gratefully acknowledge Cedric Dupuy for animal care. This work benefited from the support of the Animal Housing and Genotyping facilities funded by INSERM and LabEX BRAIN ANR-10-LABX-43. Acquisitions with the confocal Leica SP5 and slide scanner were done in the Bordeaux Imaging Center (BIC), a service unit of the CNRS-INSERM and Bordeaux University, member of the national infrastructure France BioImaging supported by the French National Research Agency (ANR-10-INBS-04). The funding bodies had no influence on design, data collection, data analysis or interpretation of data.

## AUTHOR CONTRIBUTIONS

E.P. and P.M. designed experiments. P.M. and E.C. carried out most of the experiments. E.P., P.M. and E.C. analyzed most of the data. A.B. and M.B performed the electrophysiological experiments and analyzed the corresponding data, M.F. helped for vibratome sectioning, F.C. was in charge of mouse breeding and EdU injections. C.H. provided the *Penk^Cre^* mouse line. E.P., P.M., E.C., A.B., D.N.A. discussed the results and wrote the manuscript.

