## Supplementary data for "Genetic labeling of embryonically-born dentate granule neurons in young mice using the *Penk^Cre^* mouse line"

#### SUPPLEMENTARY FIGURE LEGENDS

**Figure S1: Repartition of Penk$^{Cre}$+ cells in the mouse brain.**

PenkCre+ cells along the septo-temporal axis (coordinates from the Bregma in white) in the mouse brain at P35. One series of 40 µm sections with 400 µm between two successive sections.

Scale bar represents 2.5 mm.

**Figure S2: Birthdating of PenkCre+ cells in the two blades of the DG.**

Quantification of the percentage of ZsGreen+/EdU cells over total ZsGreen+ cells in the (A) supra and (B) infra-pyramidal blades at P35 and after injection of EdU at different time points during pregnancy, n = 3 mice per time point from at least two different litters.

**Figure S3: Contributions of principal components and synaptic properties of DGNs in *Penk$^{Cre}$;Ai6* mice**

(A) Oblique contrast image of DGNs in the suprapyramidal blade from an acute horizontal slice. A whole-cell patch-clamp pipette can be seen in contact with a ZsGreen– neuron (arrow). (B) Examples of voltage-clamp recordings from ZsGreen+ and ZsGreen– DGNs showing post synaptic currents (PSC) in the absence of pharmacological isolation. (C) (D) Boxplots of PSC frequency (Mann-Whitney test) and PSC amplitude (t-test) from ZsGreen+ and ZsGreen– neurons show no difference in post-synaptic properties. (E) Variable correlation plot showing the contribution of the selected morpho-electric parameters to the first two principal components. Parameters are color coded for their contribution.

**Figure S4: Temporal DG of *Penk$^{Cre}$; Ai6* mice at 1, 3 and 6 months**

Scale represents 50 µm.

**Figure S5: *Penk* is progressively expressed from the outer GCL to the inner GCL with age.**

(A) DG of *Penk*$^{Cre}$*;Ai14* mice in 1 month-old and 1 year-old animals. (B) *Penk* in situ hybridization in mouse sagittal sections from the Allen Brain Atlas.

Scale bars represent 100 µm (A, B)

**Figure S1**

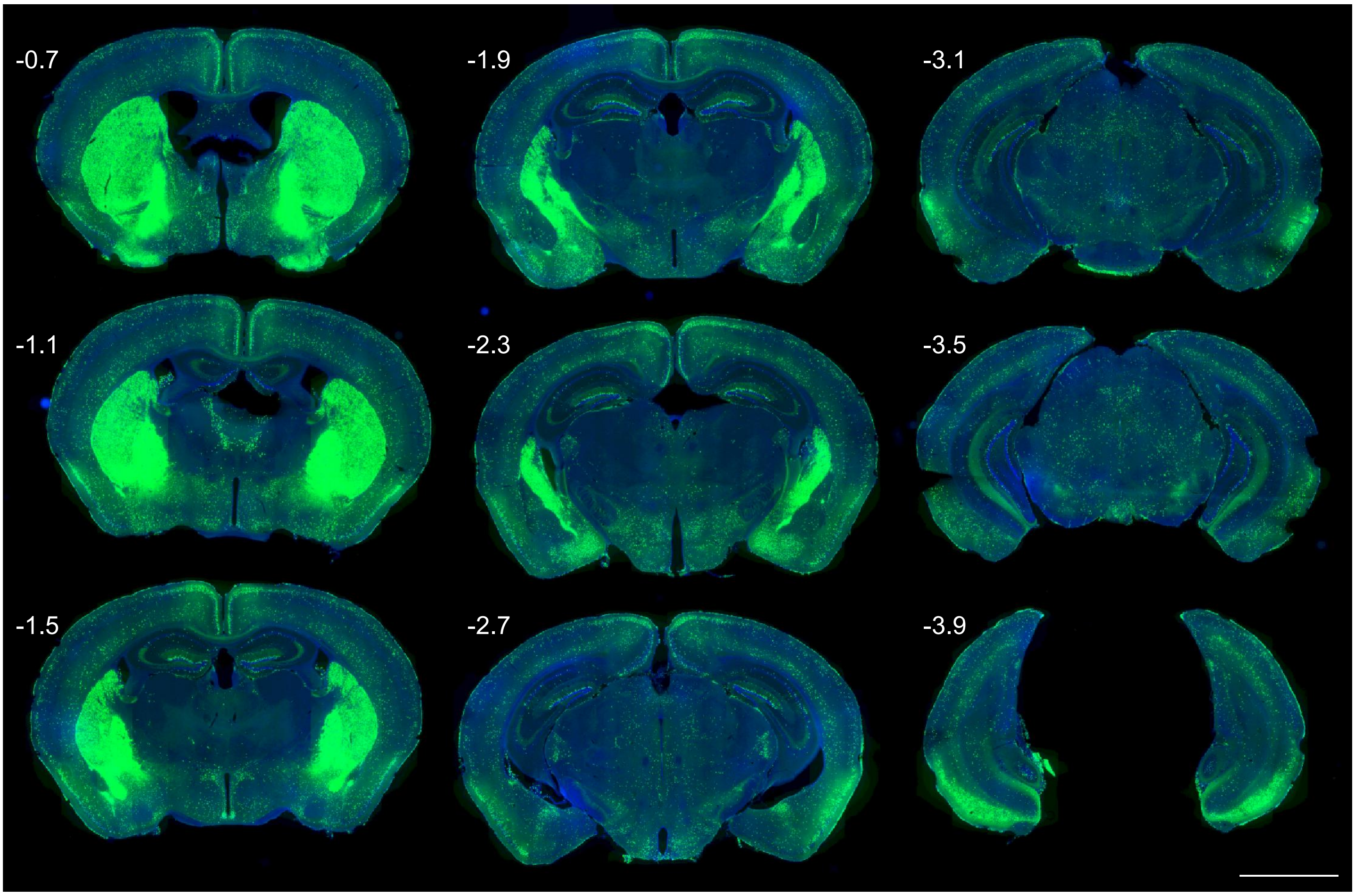

### Figure S2

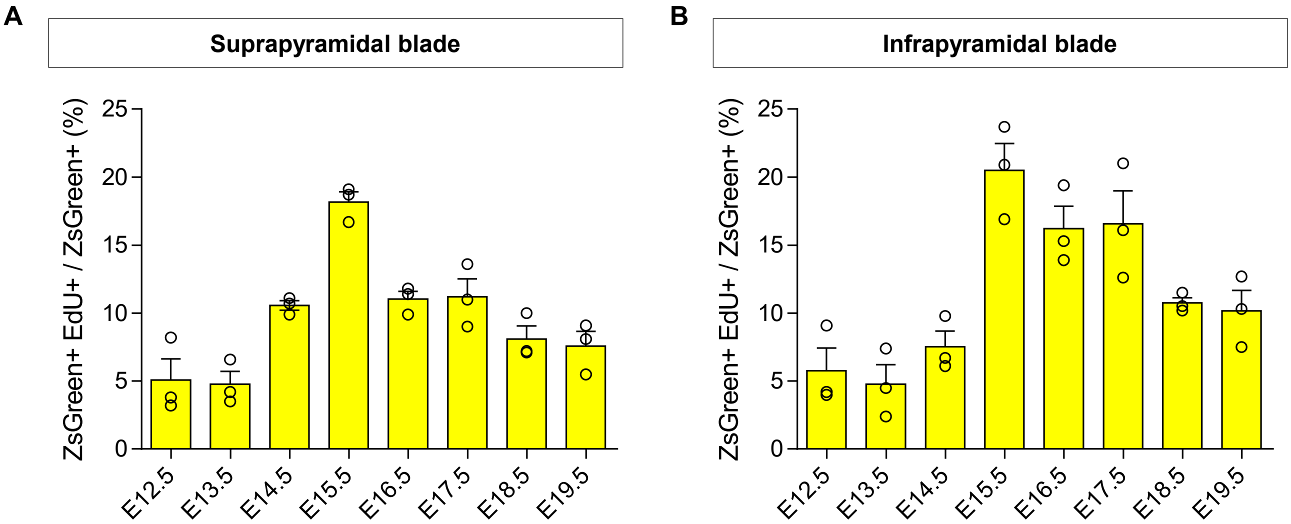

A
Suprapyramidal blade
ZsGreen+ EdU+ / ZsGreen+ (%)
0
5
10
15
20
25
E12.5
E13.5
E14.5
E15.5
E16.5
E17.5
E18.5
E19.5
B
Infrapyramidal blade
ZsGreen+ EdU+ / ZsGreen+ (%)
0
5
10
15
20
25
E12.5
E13.5
E14.5
E15.5
E16.5
E17.5
E18.5
E19.5

### Figure S3

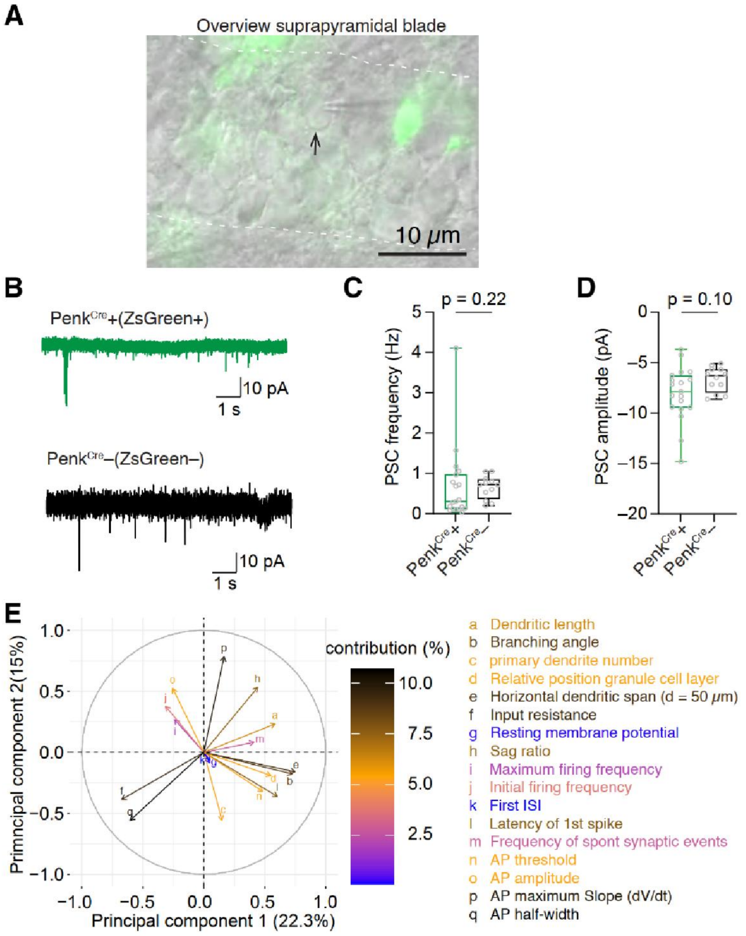

A
Overview suprapyramidal blade
10 μm
B
PenkCre+(ZsGreen+)
10 pA
1 s
PenkCre–(ZsGreen–)
10 pA
1 s
C
p = 0.22
PSC frequency (Hz)
PenkCre+
PenkCre–
D
p = 0.10
PSC amplitude (pA)
PenkCre+
PenkCre–
E
Primncipal component 2(15%)
Principal component 1 (22.3%)
contribution (%)
a Dendritic length
b Branching angle
c primary dendrite number
d Relative position granule cell layer
e Horizontal dendritic span (d = 50 μm)
f Input resistance
g Resting membrane potential
h Sag ratio
i Maximum firing frequency
j Initial firing frequency
k First ISI
l Latency of 1st spike
m Frequency of spont synaptic events
n AP threshold
o AP amplitude
p AP maximum Slope (dV/dt)
q AP half-width

### Figure S4

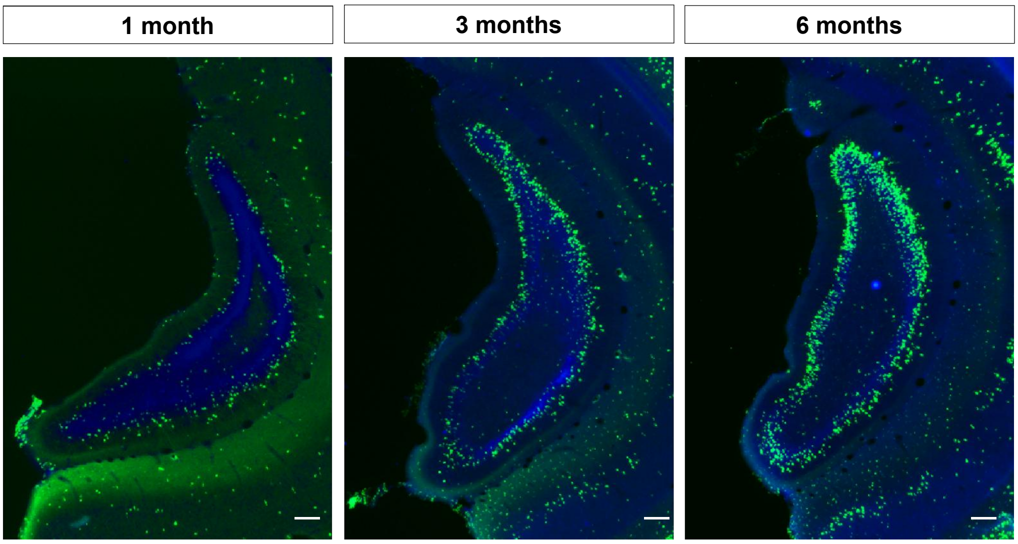

### Figure S5

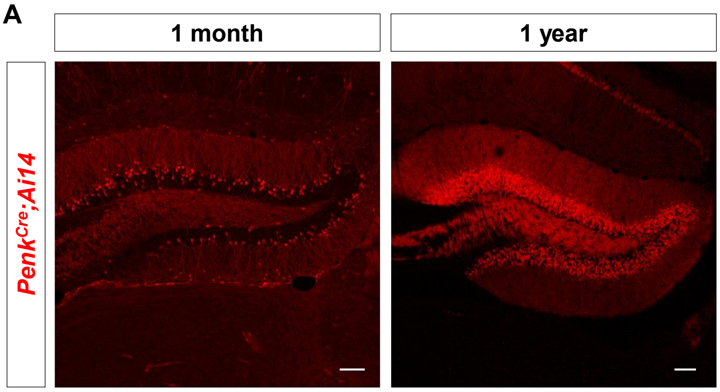
A
1 month
1 year
Penk$^{Cre}$;Ai14

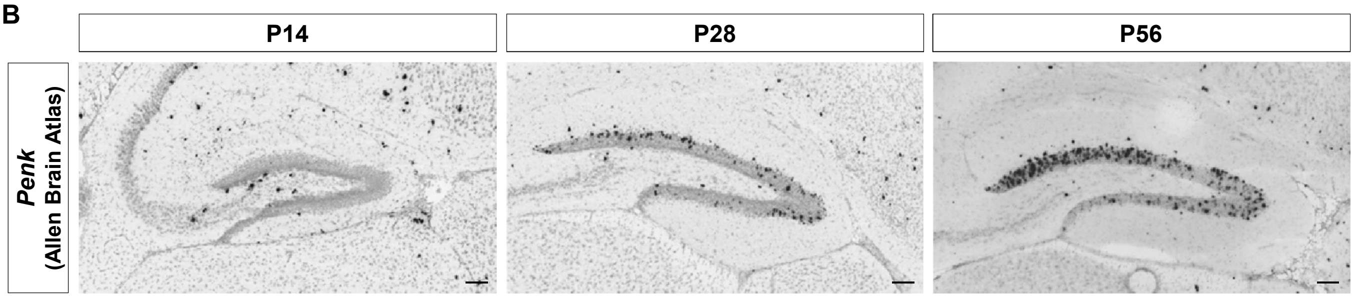
B
P14
P28
P56
Penk
(Allen Brain Atlas)
